# Muscle fiber hypertrophy in response to 6 weeks of high-volume resistance training in trained young men is largely attributed to sarcoplasmic hypertrophy

**DOI:** 10.1101/596049

**Authors:** Cody T. Haun, Christopher G. Vann, Shelby C. Osburn, Petey W. Mumford, Paul A. Roberson, Matthew A. Romero, Carlton D. Fox, Christopher A. Johnson, Hailey A. Parry, Andreas N. Kavazis, Jordan R. Moon, Veera L.D. Badsia, Benjamin M. Mwashote, Victor Ibeanusi, Kaelin C. Young, Michael D. Roberts

## Abstract

Cellular adaptations that occur during skeletal muscle hypertrophy in response to high-volume resistance training are not well-characterized. Therefore, we sought to explore how actin, myosin, sarcoplasmic protein, mitochondrial, and glycogen concentrations were altered in individuals that exhibited mean skeletal muscle fiber cross-sectional area (fCSA) hypertrophy following 6 weeks of high-volume resistance training. Thirty-one previously resistance-trained, college-aged males (mean ± standard deviation: 21±2 years, 5±3 training years) had vastus lateralis (VL) muscle biopsies obtained prior to training (PRE), at week 3 (W3), and at week 6 (W6). Muscle tissue from 15 subjects exhibiting PRE to W6 VL mean fCSA increases ranging from 320-1600 μm^2^ was further interrogated using various biochemical and histological assays as well as proteomic analysis. Seven of these individuals donated a VL biopsy after refraining from training 8 days following the last training session (W7) to determine how deloading affected biomarkers. The 15 fCSA hypertrophic responders experienced a +23% increase in mean fCSA from PRE to W6 (p<0.001) and, while muscle glycogen concentrations remained unaltered, citrate synthase activity levels decreased by 24% (p<0.001) suggesting mitochondrial volume decreased. Interestingly, both myosin and actin concentrations decreased ~30% from PRE to W6 (p<0.05). Phalloidin-actin staining similarly revealed actin concentrations per fiber decreased from PRE to W6. Proteomic analysis of the sarcoplasmic fraction from PRE to W6 indicated 40 proteins were up-regulated (p<0.05), KEGG analysis indicated that the glycolysis/gluconeogenesis pathway was upregulated (FDR sig. <0.001), and DAVID indicated that the following functionally-annotated pathways were upregulated (FDR value <0.05): a) glycolysis (8 proteins), b) acetylation (23 proteins), c) gluconeogenesis (5 proteins) and d) cytoplasm (20 proteins). At W7, sarcoplasmic protein concentrations remained higher than PRE (+66%, p<0.05), and both actin and myosin concentrations remained lower than PRE (~−50%, p<0.05). These data suggest that short-term high-volume resistance training may: a) reduce muscle fiber actin and myosin protein concentrations in spite of increasing fCSA, and b) promote sarcoplasmic expansion coincident with a coordinated up-regulation of sarcoplasmic proteins involved in glycolysis and other metabolic processes related to ATP generation. Interestingly, these effects seem to persist up to 8 days following training.

## INTRODUCTION

Weeks to months of resistance training increases skeletal muscle mean fiber cross-sectional area (fCSA) [1-10]. Tracer studies have also demonstrated that a single bout of resistance exercise increases muscle protein synthesis (MPS) and myofibrillar protein synthesis (MyoPS) rates up to 72 hours post-exercise (reviewed in [11-13]). The ingestible deuterium oxide tracer has enabled scientists to measure integrated MPS and MyoPS for prolonged periods [14, 15], and results have suggested that rates are also elevated weeks into training. Such findings have led to a general consensus that resistance training-induced skeletal muscle hypertrophy occurs via: a) increased myofibrillar protein accretion at the cellular level, and b) an increase in muscle fiber size and diameter due to said protein accretion.

While the aforementioned model is logical, there are human studies which challenge this model. For instance, a study which examined muscle fiber ultrastructural alterations in humans in response to isotonic, isometric, and run training reported decreases in the number of myofibrils per muscle fiber area across all training conditions as assessed by transmission electron microscopy (TEM) [16]. Another study [17] similarly reported significant reductions in myofibrillar area as well as an increase in sarcoplasmic area in previously untrained humans following six months of resistance training as determined through TEM imaging despite significant increases in fCSA. A similar study [18] which also utilized TEM reported that 18 weeks of resistance training resulted in a ~15% decrease in myofibrillar area (p<0.05) in both heart failure patients and healthy controls. Our laboratory recently reported that untrained college-aged males with high pre-training myofibrillar protein concentrations, as assessed through biochemical methods, experienced the largest decrease in myofibrillar protein concentrations following 12 weeks of resistance training [19]. Hence, these findings collectively suggest that the hypertrophic response to resistance training may include fiber growth through sarcoplasmic expansion (e.g., intracellular fluid, sarcoplasmic proteins, and glycogen) prior to the accretion of contractile proteins. Upon a comprehensive review of the literature, we recently defined sarcoplasmic hypertrophy as “…*an increase in the volume of the sarcolemma and/or sarcoplasm accompanied by an increase in the volume of mitochondria, sarcoplasmic reticulum, t-tubules, and/or sarcoplasmic enzyme or substrate content*” [20]. However, we also noted a severe lack of research into this area and persuaded further exploration.

We recently published a study in which 31 well-trained college-aged males completed 6 weeks of high-volume resistance training under different nutritional paradigms [21]. In light of the above, the purpose of the current study was to delineate how this training program affected actin and myosin protein concentrations, sarcoplasmic protein concentrations, glycogen concentrations, and mitochondrial volume in 15 individuals which exhibited notable increases in mean fCSA. Per the supporting literature above, we hypothesized that individuals experiencing notable fCSA increases would experience a decrease in myosin and actin concentrations, a decrease in citrate synthase activity (indicative of decreased mitochondrial content), and either no change or an increase in sarcoplasmic protein and glycogen concentrations.

## MATERIALS AND METHODS

### Ethical approval and study design

Prior to engaging in data collection, this study was approved by the Institutional Review Board at Auburn University (Protocol #17-425 MR 1710; IRB contact) and conformed to the standards set by the latest revision of the Declaration of Helsinki. Resistance-trained college-aged males from the local community were recruited to participate in this study. Participants provided both verbal and written informed consent, and completed a medical history form prior to screening. Two primary criteria were used to establish training status: a) greater than one year of resistance training (self-reported), and b) a back squat one repetition maximum (1RM) ≥1.5× body mass.

### Establishment of mean fCSA responders

We recently analyzed serial muscle sections from 26 participants and reported that the standard error of measurement for detecting fCSA changes in our laboratory is ±459 μm^2^ [20]. Out of the 31 participants only 11 experienced >459 μm^2^ increase in mean fCSA. However, in order to increase statistical power and considering the exploratory nature of this work, we analyzed specimens from a total of 15 subjects demonstrating notably positive PRE to W6 mean fCSA increases ranging from 320-1600 μm^2^. In the original study, participants were partitioned to one of three nutritional supplement groups including: a) daily single dose of whey protein (WP, 25g/d; n=10), b) daily single dose of maltodextrin (MALTO, 30 g/d; n=10), or c) graded dose of WP (GWP, 25-150 g/d from weeks 1-6; n=11). In the 15 participants experiencing mean FCA increases >320 μm^2^ there was practically equal representation of participants from each of the supplement groups (MALTO n=6, WP n=4, GWP n=5) suggesting that results were likely not influenced from nutritional supplementation. Readers are referred to Haun et al. [21] for more in-depth descriptions of the supplementation details and nutritional recommendations employed.

### PRE, W3, and W6 testing sessions

Prior to training (PRE), following week 3 (W3), and following week 6 (W6) participants arrived for testing batteries in an overnight fasted condition. At W3 and W6, testing sessions occurred 24 hours following Friday workouts. Testing procedures pertinent to this dataset are described below, although other assessments were performed and described in greater detail elsewhere [21].

#### Urine specific gravity testing for adequate hydration

At the beginning of each testing session, participants submitted a urine sample (~5 mL) to assess urine specific gravity (USG) levels (1.005-1.020 ppm) using a handheld refractometer (ATAGO; Bellevue, WA, USA) as a measure of hydration status. Participants with a USG ≥1.020 were asked to consume 400 mL tap water and were re-tested 20 minutes thereafter.

#### Muscle tissue and blood collection

Muscle biopsies were collected using a 5-gauge needle under local anesthesia as previously described [21]. Immediately following tissue procurement, ~20-40 mg of tissue was embedded in cryomolds containing optimal cutting temperature (OCT) media (Tissue-Tek®, Sakura Finetek Inc; Torrence, CA, USA). Embedding was performed according to methods previously published by our laboratory [22] whereby tissue was placed in cryomolds for cross sectional slicing in a non-stretched state prior to rapid freezing. Cryomolds were then frozen using liquid nitrogen-cooled isopentane and subsequently stored at −80°C until histological staining for determination of mean fCSA and actin protein per fiber as described below. The remaining tissue was teased of blood and connective tissue, wrapped in pre-labelled foils, flash frozen in liquid nitrogen, and subsequently stored at −80°C until molecular analyses were performed as described below. Venous blood samples were also collected into a 5 mL serum separator tube (BD Vacutainer, Franklin Lakes, NJ, USA) during the waiting period for anesthesia to take effect for creatine kinase (CK) activity assessment described in greater detail below.

### Resistance training protocol

Participants were familiarized with the design of training and technical parameters during testing of 3RMs which occurred 3-7 days prior to PRE testing and training initiation. Strict technical parameters were employed for testing to ensure accurate reflections of strength under direct supervision of research staff holding the Certified Strength and Conditioning Specialist Certification from the National Strength and Conditioning Association.

Following the PRE testing battery and 3RM testing, RT occurred 3 days per week. Loads corresponding to 60% 1RM, based on 3RM testing, were programmed for each set of each exercise. Sets of 10 repetitions were programmed for each set of each exercise throughout the study. Exercises were completed one set at a time, in the following order during each training session: days 1 and 3 each week – barbell (BB) back squat, BB bench press, BB stiff-legged deadlift, and an underhand grip cable machine pulldown exercise designed to target the elbow flexors and latissimus dorsi muscles (lat pulldown); day 2 of each week – BB back squat, BB overhead press, BB stiff-legged deadlift, and lat pulldown. The 3 day per week protocol involved a progressive increase from 10 sets per week to 32 sets per week for each exercise. Thus, on the last week of training participants performed 32 sets of 10 repetitions of BB back squats, 32 sets of 10 repetitions of BB bench press and overhead press combined, 32 sets of 10 repetitions of BB stiff-legged deadlift, and 32 sets of 10 repetitions of lat pulldowns. Readers are referred to Haun et al. [21] for more in-depth descriptions of training.

### Molecular analyses

Pulverized tissue was used for Western blotting, citrate synthase activity, 20S proteasome activity, and glycogen concentration assays. Tissue foils were removed from −80°C and tissue was crushed on a liquid nitrogen-cooled ceramic mortar with a ceramic pestle. Powdered tissue was weighed using an analytical scale sensitive to 0.0001 g (Mettler-Toledo; Columbus, OH, USA), and placed in 1.7 mL polypropylene tubes containing 500 μL of ice-cold general cell lysis buffer [20 mM Tris-HCl (pH 7.5), 150 mM NaCl, 1 mM Na_2_EDTA, 1 mM EGTA, 1% Triton; Cell Signaling, Danvers, MA, USA] pre-stocked with protease and Tyr/Ser/Thr phosphatase inhibitors (2.5 mM sodium pyrophosphate, 1 mM β-glycerophosphate, 1 mM Na_3_VO_4_, 1 μg/mL leupeptin). Samples were then homogenized by hand using tight-fitting micropestles, insoluble proteins were removed via centrifugation at 500 *g* for 5 minutes, and supernatants (whole-tissue sample lysates) were stored at −80°C and used for further molecular analyses described below.

#### Protein concentration determination

Whole-tissue sample lysates were batch process-assayed for total protein content using a BCA Protein Assay Kit (Thermo Fisher Scientific; Waltham, MA, USA). These lysates were used for Western blotting and enzyme assays described below.

#### Western blotting

Lysates were then prepared for Western blotting using 4x Laemmli buffer at 1 μg/μL and boiled for 5 minutes at 100°C. Samples (15 μL) were loaded onto pre-casted gradient (4-15%) SDS-polyacrylamide gels (Bio-Rad Laboratories; Hercules, CA, USA) and subjected to electrophoresis at 180 V for 60 minutes using pre-made 1x SDS-PAGE running buffer (Ameresco; Framingham, MA, USA). Proteins were then transferred (200 mA for 2 hours) to polyvinylidene difluoride membranes (Bio-Rad Laboratories), Ponceau stained, and imaged to ensure equal protein loading between lanes. Membranes were then blocked for 60 minutes at room temperature with 5% nonfat milk powder in Tris-buffered saline with 0.1% Tween-20 (TBST; Ameresco). Rabbit anti-human ubiquitin (1:1,000 dilution; catalog #: 3933; Cell Signaling) in TBST with 5% bovine serum albumin (BSA) was added to membranes and incubated overnight at 4°C. The following day, membranes were washed and incubated with horseradish peroxidase-conjugated anti-rabbit IgG (catalog #: 7074; Cell Signaling) in TBST (1:2,000 dilution) with 5% BSA at room temperature for 1 hour. Membranes were developed using an enhanced chemiluminescent reagent (Luminata Forte HRP substrate; EMD Millipore, Billerica, MA, USA), and band densitometry was performed using a gel documentation system with associated software (UVP, Upland, CA, USA). Whole-lane densitometry values for poly-ubiquinated proteins of each subject at each time point were divided by whole-lane Ponceau densities in order to derive relative expression levels.

#### Citrate synthase activity assay

Whole-tissue sample lysates were batch processed for citrate synthase activity as previously described [19], and this marker was used as a surrogate for mitochondrial content according to previous literature suggesting citrate synthase activity highly correlates with electron micrograph images of mitochondrial content in skeletal muscle (r=0.84, p<0.001) [23]. The assay utilized is based on the reduction of 5, 50-dithiobis (2-nitrobenzoic acid) (DTNB) at 412 nm (extinction coefficient 13.6 mmol/L/cm) coupled to the reduction of acetyl-CoA by the citrate synthase reaction in the presence of oxaloacetate. Briefly, 12.5 μg of whole-tissue sample lysates were added to a mixture composed of 0.125 mol/L Tris-HCl (pH 8.0), 0.03 mmol/L acetyl-CoA, and 0.1 mmol/L DTNB. The reaction was initiated by the addition of 5 μL of 50 mM oxaloacetate and the absorbance change was recorded for one minute. The average coefficient of variation values for all duplicates was 5.2%.

#### 20S proteasome activity assay

Protein from whole-tissue sample lysates (40 μg) were batch processed for 20S proteasome activity using commercially available fluorometric kits (catalog #: APT280; Millipore Sigma; Burlington, MA, USA) per the manufacturer’s instructions. Assay readings are presented as raw fluorometric values as previously described by our laboratory [24]. The average coefficient of variation values for all duplicates was 8.7%.

#### Glycogen assay

Whole-tissue sample lysates were batch processed for glycogen determination using commercially-available fluorometric kits (catalog #: MAK016; Millipore Sigma) as per the manufacturer’s instructions. This assay was piloted with whole-tissue sample lysates, frozen wet tissue, and lyophilized tissue, and all three methods yielded similar quantitative results. Thus, given that whole-tissue sample lysates were available for most participants, we opted to assay this tissue fraction. A standard curve was used to determine glycogen content of whole-tissue sample lysates, this value was multiplied by the starting cell lysis buffer volume (500 μL) to derive total glycogen content per sample, and the resultant value was divided by input wet muscle weights to obtain nmol glycogen/mg wet muscle weight. The average coefficient of variation values for all duplicates was 9.8%.

### Assessment of sarcoplasmic protein, myosin, and actin

Myofibrillar and sarcoplasmic protein isolations were performed based on the methods of Goldberg’s laboratory [25]. Briefly, frozen wet powdered muscle (~8-20 mg) was weighed, placed into 1.7 mL polypropylene tubes with ventilation holes drilled on the caps. Samples were freeze-dried using a laboratory-grade freeze-drying device (FreeZone 2.5, Labconco; Kansas City, MO, USA). Freeze dry settings were −50°C for the condenser and a vacuum setting of 0.1 mBar, and samples were placed in the condenser well for 4 hours under vacuum pressure. Following freeze drying, muscle was re-weighed on an analytical scale to determine muscle fluid content, and placed in 1.7 mL polypropylene tubes containing 190 μL of ice cold homogenizing buffer (20 mM Tris-HCl, pH 7.2, 5 mM EGTA, 100 mM KCl, 1% Triton-X 100). Samples were homogenized on ice using tight-fitting pestles, and centrifuged at 3000 *g* for 30 minutes at 4°C. Supernatants (sarcoplasmic fraction) were collected, placed in 1.7 mL polypropylene tubes, and stored at −80°C. The resultant myofibril pellet was resuspended in homogenizing buffer, and samples were centrifuged at 3000 *g* for 10 minutes at 4°C. Resultant supernatants from this step were discarded, pellets were resuspended in 190 μL of ice cold wash buffer (20 mM Tris-HCl, pH 7.2, 100 mM KCl, 1 mM DTT), and samples were centrifuged at 3000 *g* for 10 minutes at 4°C; this specific process was performed twice and supernatants were discarded during each step. Final myofibril pellets were resuspended in 200 μL of ice cold storage buffer (20 mM Tris-HCl, pH 7.2, 100 mM KCl, 20% glycerol, 1 mM DTT) and frozen at −80°C. Sarcoplasmic protein content was determined in duplicate using a cuvette-based BCA assay protocol (Thermo Fisher Scientific) whereby a large volume of sample was assayed (100 μL of 5× diluted sample + 900 μL Reagent A + B). The average CV for duplicate readings was 0.58%, and sarcoplasmic protein concentrations were normalized to input dry muscle weights.

#### Electrophoretic determination of myosin and actin content

Determination of myosin and actin content were performed as previously described by our laboratory [19] and others [25]. Briefly, SDS-PAGE preps were made using 10 μL resuspended myofibrils, 65 μL distilled water (diH_2_O), and 25 μL 4x Laemmli buffer. Samples (15 μL) were then loaded on pre-casted gradient (4-15%) SDS-polyacrylamide gels (Bio-Rad Laboratories) and subjected to electrophoresis at 200 V for 40 minutes using pre-made 1x SDS-PAGE running buffer (Ameresco). Following electrophoresis gels were rinsed in diH_2_O for 15 minutes, and immersed in Coomassie stain (LabSafe GEL Blue; G-Biosciences; St. Louis, MO, USA) for 2 hours. Thereafter, gels were destained in diH_2_O for 60 minutes, bright field imaged using a gel documentation system (UVP), and band densities were determined using associated software. Given that a standardized volume from all samples were loaded onto gels, myosin and actin band densities were normalized to input muscle weights for relative expression. Our laboratory has reported that this method yields exceptional sensitivity in detecting 5-25% increases in actin and myosin content [19]. Notably, actin and myosin content were the only two myofibrillar protein targets of interest given that the combination of these proteins make up a majority (~70%) of the myofibrillar protein pool [20].

#### Proteomic analysis of the sarcoplasmic fractions from PRE and W6

Proteomics analysis was performed on the sarcoplasmic fraction for n=12 fCSA responders in order to interrogate the sarcoplasmic proteins which were altered from PRE to W6. Each protein (90 μg) sample for triplicate technical runs (30 μg for each run) was prepared for LC-MS/MS analysis using EasyPep™ Mini MS Sample Prep Kit (Thermo Fisher Scientific). In brief, protein sample was transferred into a new microcentrifuge tube and the final volume was adjusted to 100 μL with general cell lysis buffer (Cell Signaling). Reduction solution (50 μL) and alkylation solution (50 μL) were added to the sample, gently mixed, and incubated at 95°C using heat block for 10 minutes to reduce and alkylate samples. After incubation, the sample was removed from the heat block and cooled to room temperature. The reconstituted enzyme Trypsin/Lys-C Protease Mix solution (50 μL) was added to the reduced and alkylated protein sample and incubated with shaking at 37°C for 2 hours to digest the protein sample. Digestion stop solution (50 μL) was added to the sample and peptides were cleaned using peptide clean-up column according to the kit instructions.

An externally calibrated Thermo Q Exactive HF (high-resolution electrospray tandem mass spectrometer) was used in conjunction with Dionex UltiMate3000 RSLC Nano System (Thermo Fisher Scientific). Sample (5 μL) was aspirated into a 50 μL loop and loaded onto the trap column (Thermo μ-Precolumn 5 mm, with nanoViper tubing 30 μm i.d. × 10 cm). The flow rate was set to 300 nL/min for separation on the analytical column (Acclaim pepmap RSLC 75 μM × 15 cm nanoviper; Thermo Fisher Scientific). Mobile phase A was composed of 99.9% H_2_O (EMD Omni Solvent; Millipore, Austin, TX, USA), and 0.1% formic acid and mobile phase B was composed of 99.9% acetonitrile and 0.1% formic acid. A 60 min linear gradient from 3% to 45% B was performed. The LC eluent was directly nanosprayed into the mass-spectrometer. During chromatographic separation, the Q-Exactive HF was operated in a data-dependent mode and under direct control of the Thermo Excalibur 3.1.66 (Thermo Fisher Scientific). The MS data were acquired using the following parameters: 20 data-dependent collisional-induced-dissociation (CID) MS/MS scans per full scan (350 to 1700 m/z) at 60,000 resolution. MS2 were acquired in centroid mode at 15,000 resolution. Ions with a single charge or charges more than 7 as well as unassigned charge were excluded. A 15-second dynamic exclusion window was used. All measurements were performed at room temperature, and three technical replicates were run for each sample. The raw files were analyzed using Proteome Discoverer (version 2.0, Thermo Fisher Scientific) software package with SequestHT and Mascot search nodes using species specific tremble fasta database and the Percolator peptide validator. The resulting.msf files were further analyzed by the proteome validator software Scaffold v4.4 (Portland, OR, USA).

### Immunohistochemistry for fCSA and phalloidin staining

Methods for cryostat sectioning have been employed previously in our laboratory and described elsewhere [21, 22, 26, 27]. Briefly, sections from OCT-preserved samples were cut at a thickness of 8 μm using a cryotome (Leica Biosystems; Buffalo Grove, IL, USA) and were adhered to positively-charged histology slides. Once all samples were sectioned, microscope slides were stored at −80°C until batch processing occurred for immunohistochemistry.

For mean fCSA determination, sections were air-dried at room temperature for 10 minutes, permeabilized in a phosphate-buffered saline (PBS) solution containing 0.5% Triton X-100, and blocked with 100% Pierce Super Blocker (Thermo Fisher Scientific) for 10 minutes. Sections were then incubated for 15 minutes with a pre-diluted commercially-available rabbit anti-dystrophin IgG antibody solution (catalog #: GTX15277; Genetex Inc.; Irvine, CA, USA) and spiked in mouse anti-myosin I IgG (catalog #: A4.951 supernatant; Hybridoma Bank, Iowa City, IA, USA; 40 μL added per 1 mL of dystrophin antibody solution). Sections were then washed for 2 minutes in PBS and incubated in the dark for 15 minutes with a secondary antibody solution containing Texas Red-conjugated anti-rabbit IgG (catalog #: TI-1000; Vector Laboratories, Burlingame, CA, USA), and Alexa Fluor 488-conjugated anti-mouse IgG (catalog #: A-11001; Thermo Fisher Scientific; 6.6 μL of all secondary antibodies per 1 mL of blocking solution). Sections were washed for 2 minutes in PBS, air-dried and mounted with fluorescent media containing 4,6-diamidino- 2-phenylindole (DAPI; catalog #: GTX16206; Genetex Inc.). Following mounting, slides were stored in the dark at 4°C until fluorescent images were obtained. After staining was performed on all sections, digital 10× objective images were captured using a fluorescence microscope (Nikon Instruments, Melville, NY, USA). All images were captured by a laboratory technician who was blinded to the labeling codes on microscope slides. Approximate exposure times were 400 ms for TRITC and FITC imaging, and 80 ms for DAPI imaging. This staining method allowed the identification of cell membranes (detected by the Texas Red filter), type I fiber green cell bodies (detected by the FITC filter), type II fiber black cell bodies (unlabeled), and myonuclei (detected by the DAPI filter). Combined CSA measurements of both fiber types (mean fCSA) were performed using custom-written pipelines in the open-sourced software CellProfiler^TM^ [28] whereby the number of pixels counted within the border of each muscle fiber was converted to a total area (μm^2^). A calibrator slide containing a 250,000 μm^2^ square image was also captured, and pixels per fiber from imaged sections were converted to area using this calibrator image. On average, 113±26 fibers per cross-section were identified for analysis at each sampling time. A post hoc experiment performed in our laboratory to examine potential differences in fCSA measurements between sections on the same slide (n=27 slides) revealed strong reliability using this method (ICC=0.929).

For actin protein content determination per fiber cross section, F-actin labelling using Alexa Fluor 488-conjugated (AF488) phalloidin was performed according to previous reports to estimate actin protein content per muscle fiber cross section [29, 30]. Briefly, sections were air-dried for 10 minutes and fixed in formalin for 10 minutes. Sections were then washed with PBS for 5 minutes, and blocked with 100% Pierce Super Blocker (Thermo Fisher Scientific) for 25 minutes. After blocking, a solution containing 5% Super Blocker, 1% rabbit anti-human collagen I IgG (catalog #: ab34710; Abcam, Cambridge, MA, USA), and 3% phalloidin-AF488 (catalog #: A12379; Thermo Fisher Scientific) was placed on sections in the dark for 30 minutes. Sections were subsequently washed for 5 minutes with PBS and incubated in the dark for 20 minutes with a secondary antibody solution containing Texas Red-conjugated anti-rabbit IgG (catalog #: TI-1000; Vector Laboratories; ~6.6 μL secondary antibody per 1 mL of blocking solution). Sections were washed for 5 minutes in PBS, air-dried and mounted with fluorescent media (Genetex Inc.). Following mounting, slides were stored in the dark at 4°C until immunofluorescent images were obtained. After staining was performed on all sections, digital images were captured from blinded samples using a fluorescence microscope (Nikon Instruments) using the 20× objective. Exposure times were 80 milliseconds for FITC and 2 seconds for TRITC imaging. This staining method allowed the identification of actin protein (detected by the FITC filter) and outer cell matrix (detected by the Texas Red filter). Automated measurements of phalloidin-stained pixel intensity per fiber were performed using CellProfiler^TM^ whereby the number of pixels counted within the border of each muscle fiber were converted summed to provide an integrated intensity value. This value was then divided by fiber area in order to derive a relative contractile intensity value per fiber per μm^2^.

### Serum analysis

Upon blood collection, serum tubes were centrifuged at 3,500 *g* for 5 minutes at room temperature. Aliquots were then placed in 1.7 mL polypropylene tubes and stored at −80°C until batch-processing. An activity assay was used to determine serum levels of CK (Bioo Scientific, Austin, TX, USA). All kits were performed according to manufacturer's instructions and plates were read using a 96-well spectrophotometer (BioTek, Winooski, VT, USA). Due to missed time points and one subject demonstrating exceptionally high PRE serum CK activity values, only 11 subjects were included in the PRE-W6 analysis, and only 6 subjects were included in the W7 analysis described below.

### W7 biopsy assays from a subset of fCSA responders

VL biopsies were obtained from subset of fCSA responders (n=7) 8 days following the last training bout (termed W7). These individuals refrained from training between W6 and W7, and the purpose of this visit was to ensure that training-induced damage had subsided following transient deloading. Due to financial constraints, only select assays were performed on these subjects and are presented in the results.

### Statistical analysis

Dependent variable comparisons over time were analyzed using one-way repeated measures ANOVAs with LSD post hoc tests. When sphericity was violated (Mauchly's Test of Sphericity p<0.05), Greenhouse-Geisser corrections were applied. Pearson correlations were also performed on select variables. All statistical analyses were performed using SPSS v22.0 (IBM Corp, Armonk, NY, USA) and significance was established at p<0.05, although p-values “approaching significance” (i.e., 0.050<p<0.100) were also discussed given the exploratory nature of the investigation. There was inadequate VL biopsy tissue for certain assays or sections for phalloidin staining. Additionally, only 12 participants were analyzed using proteomics analysis due to financial constraints. Thus, participant number for each dependent variable are presented in figures.

## RESULTS

### PRE, W3, and W6 alterations in VL biopsy molecular variables in fCSA responders

Figure 1a and b illustrates the 15 hypertrophic responders according to fCSA changes from PRE to W6. There were no significant changes in VL biopsy glycogen concentrations (Figure 1c), fluid content (Figure 1d), or sarcoplasmic protein concentrations (Figure 1f). Citrate synthase activity decreased from PRE to W3 (p<0.001), and activity also remained lower than PRE at W6 (p<0.001) (Figure 1e). Interestingly, myosin and actin protein concentrations significantly decreased from PRE to W6 (p=0.035 for each target; Figure 1g-i). Figure 1j demonstrates that the utilized myofibrillar and sarcoplasmic isolation methods show excellent enrichment of contractile proteins in the former fraction, and little contamination of contractile proteins in the latter fraction.

**Figure 1.**
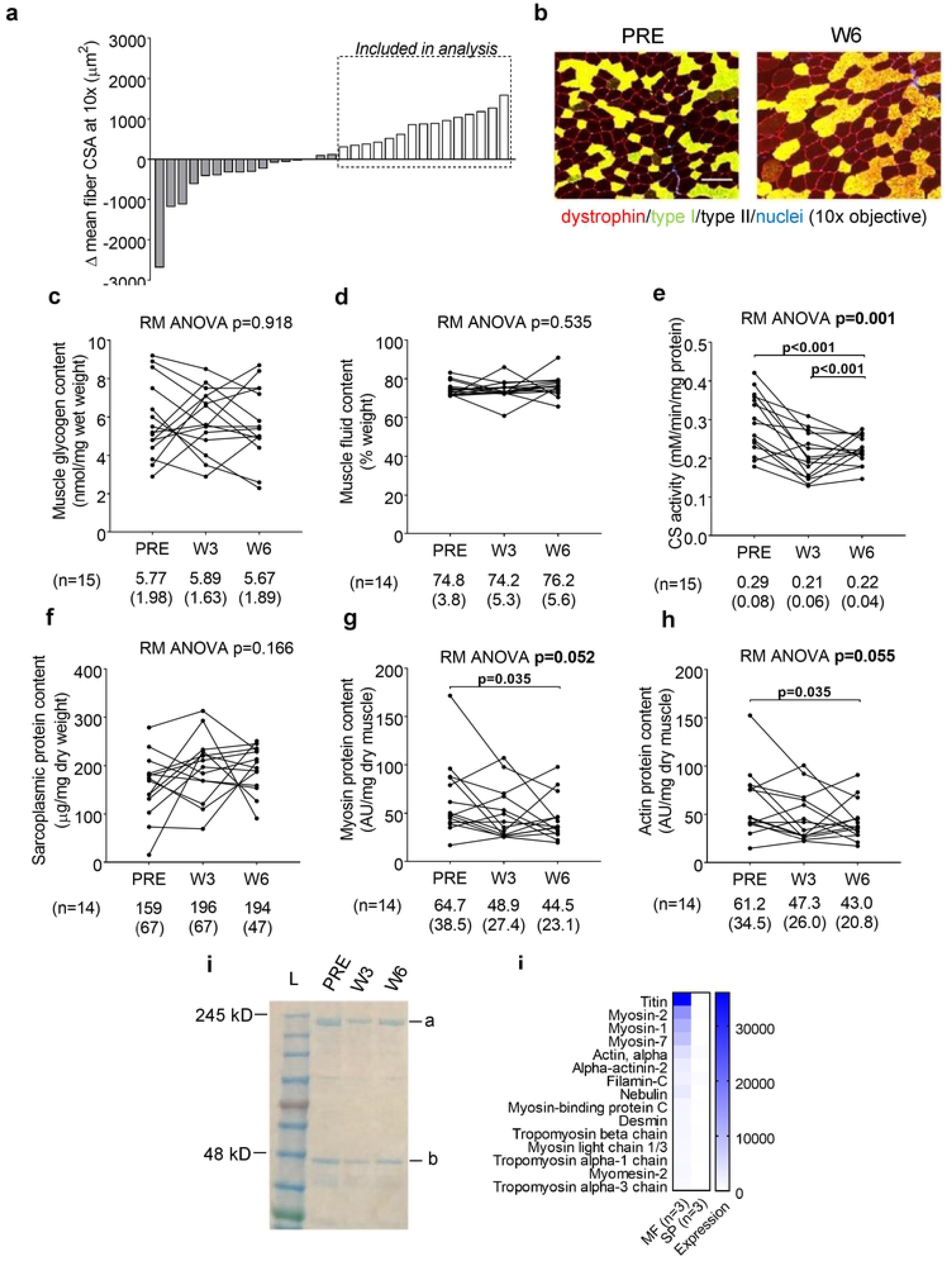
Right leg body composition and VL biopsy molecular adaptations in fCSA responders. Legend: All line graph data show individual responses with mean ± standard deviation (SD) values below each time point. Panel a demonstrates the 15 hypertrophic responders according to fCSA changes with training. Panel b illustrates a PRE and W6 representative 10× objective image from one subject (scale bar = 200 μm). No significant changes in VL biopsy glycogen concentrations (panel c), muscle fluid content (panel d), or sarcoplasmic protein concentrations (panel f) were observed. VL biopsy citrate synthase activity (a surrogate of mitochondrial volume) significantly decreased over time (panel e). VL biopsy myosin protein (panel g) and actin protein (panel h) concentrations significantly decreased from PRE to W6. Panel i illustrates representative myosin (band a) and actin (band b) Coomassie stains from one subject. Panel j demonstrates that the utilized myofibrillar and sarcoplasmic isolation methods demonstrated excellent enrichment of contractile proteins in the former fraction, and little contamination of contractile proteins in the latter fraction (both fractions from n=3 random participants were assayed).

### PRE, W3, and W6 VL biopsy actin content per muscle fiber assessed through phalloidin-AF488 staining in fCSA responders

Figure 2 demonstrates phalloidin-AF488 staining results of 11 of the 15 subjects who yielded muscle tissue of adequate quality for staining. Phalloidin integrated pixel intensity (PIPI) per fiber represents the amount of actin per fiber cross-section, and an increase in this metric with training indicates muscle fiber actin accretion. Interestingly, PIPI was not significantly altered in these 11 fCSA responders throughout the intervention indicating there was no significant actin accretion (Figure 2a). Phalloidin pixel intensity per fiber per μm^2^ represents contractile protein density (or concentration), and no changes in this metric would indicate that actin accretion occurs in proportion to muscle fiber growth whereas a decrease in this metric would indicate a reduction in actin concentration. The data in Figure 2b represent the W3 and W6 change scores in this metric from the PRE time point. There were numerical but non-significant decreases from PRE to W3 (p=0.226) and W6 (p=0.519). However, the removal of subject 17 (statistical outlier >2 SD from mean) indicated a significant decrease occurred from PRE to W6 (p=0.020). Another insightful analysis included examining the association between fCSA and PIPI, where a very high association in this variable indicates the coupling of fiber size and actin content. There was tight coupling at PRE (r^2^=0.815, p<0.001; Figure 2c) and W3 (r^2^=0.867, p<0.001; Figure 2d) suggesting a proportional relationship existed between fCSA and actin content per fiber at these time points. However, a weak non-significant correlation was observed at W6 (r^2^=0.160, p=0.222; Figure 2e) suggesting a disproportionate increase in fCSA occurred without increases in actin content per fiber. Figure 2f demonstrates a representative phalloidin-AF488 staining image from a participant at the PRE and W6 time points.

**Figure 2.**
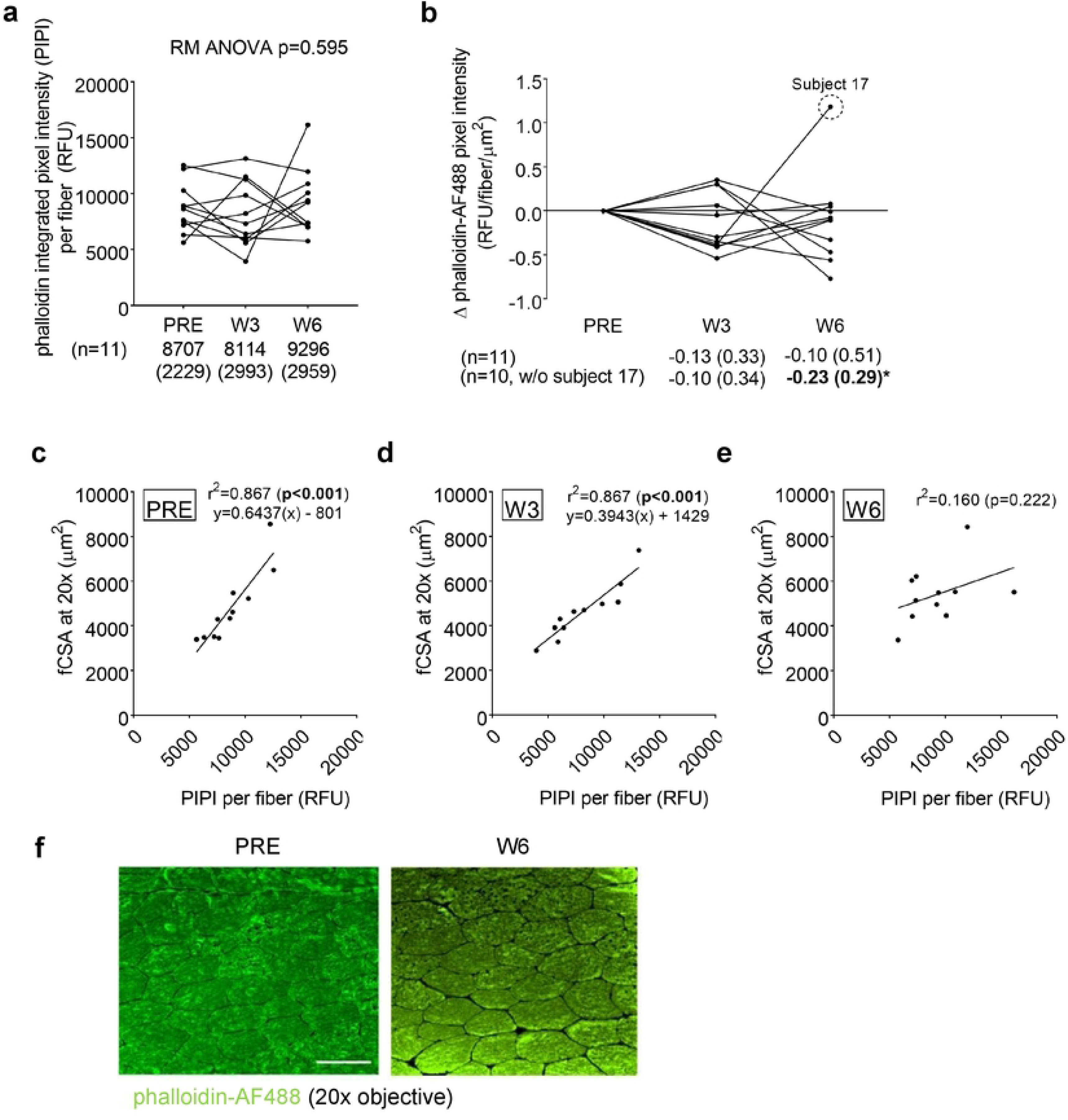
PRE, W3, and W6 VL biopsy contractile protein per muscle fiber assessed through phalloidin-AF488 staining. Legend: All line graph data show individual responses with mean ± standard deviation (SD) values below each time point. Phalloidin-AF488 integrated pixel intensity (PIPI) per fiber represents the amount of actin per fiber where increases in this metric suggest increases in the amount of actin per muscle fiber. This metric was not significantly altered throughout the intervention (panel a). Phalloidin pixel intensity per fiber per μm^2^, which represents actin content corrected for fCSA, demonstrated numerical but non-significant decreases from PRE to W3 and W6 (panel b), although the removal of subject 17 (statistical outlier) indicated a significant decrease from occurred from PRE to W6. Panels c and d demonstrate a tight coupling at PRE and W3 between fCSA and contractile protein content per fiber. However, only a modest and non-significant correlation was observed at W6 (panel e). Panel g illustrates PRE and W6 phalloidin-AF488 stain 20× images from one subject (scale bar = 100 μm).

### PRE, W3, and W6 VL biopsy and serum markers of proteolysis and damage in fCSA responders

Given that actin and myosin concentrations decreased from PRE to W6, we were interested in examining if markers of proteolysis or damage were concomitantly elevated. However, 20S proteasome activity, ubiquinated protein levels, and serum CK activity did not significantly change over time (Figures 3a-c). Figure 3d demonstrates a representative Western blot image of ubiquinated protein levels from a participant at the PRE, W3, and W6 time points.

**Figure 3.**
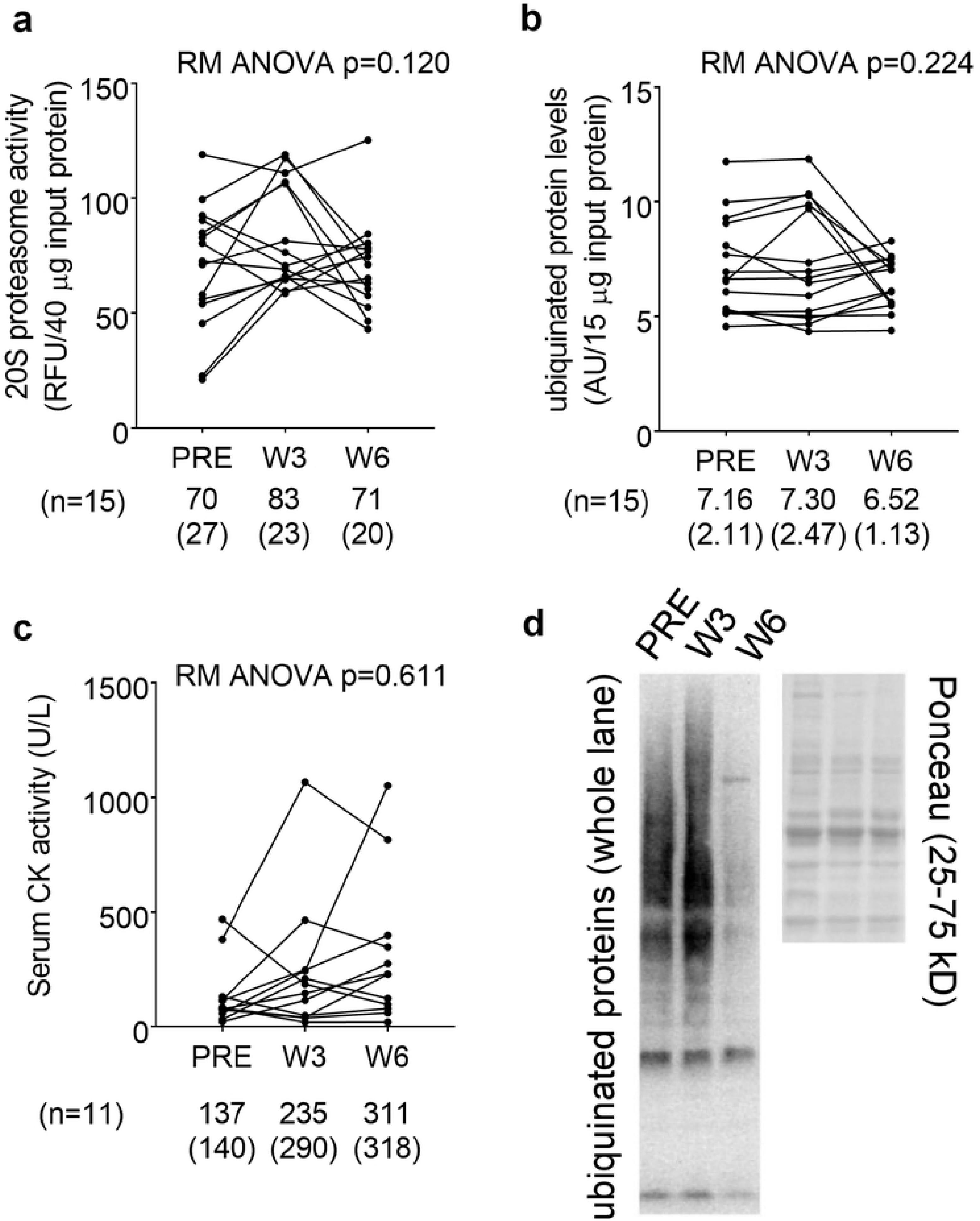
PRE, W3, and W6 VL biopsy proteolysis and muscle damage markers. Legend: All line graph data show individual responses with mean ± standard deviation (SD) values below each time point. 20S proteasome activity (panel a), ubiquinated protein levels (panel b), and serum CK activity (panel c) did not significantly change over time. Panel d illustrates a representative Western blot image of ubiquinated protein levels from one subject.

### Proteomics analysis of the PRE and W6 sarcoplasmic fractions from fCSA responders

Given that sarcoplasmic protein concentrations numerically increased from PRE to W6 and actin and myosin protein concentrations concomitantly decreased, we sought to examine which sarcoplasmic proteins were altered. Of the 157 sarcoplasmic proteins detected, 40 proteins were significantly upregulated (p<0.05; Figure 4a), and 1 protein was significantly down-regulated from PRE to W6 (tropomyosin beta chain, 0.29-fold relative to PRE; *data not shown on* Figure 4). Kyoto Encyclopedia of Genes and Genomes (KEGG) pathway analysis [31] indicated that only one pathway termed glycolysis/gluconeogenesis (8 up-regulated proteins) was significantly up-regulated from PRE to W6 (FDR value < 0.001; Figure 4b). Pathway analysis using DAVID v6.8 [32] indicated that the following functionally-annotated pathways were significantly upregulated (FDR value <0.05): a) glycolysis (8 proteins), b) acetylation (23 proteins), c) gluconeogenesis (5 proteins) and d) cytoplasm (20 proteins).

**Figure 4.**
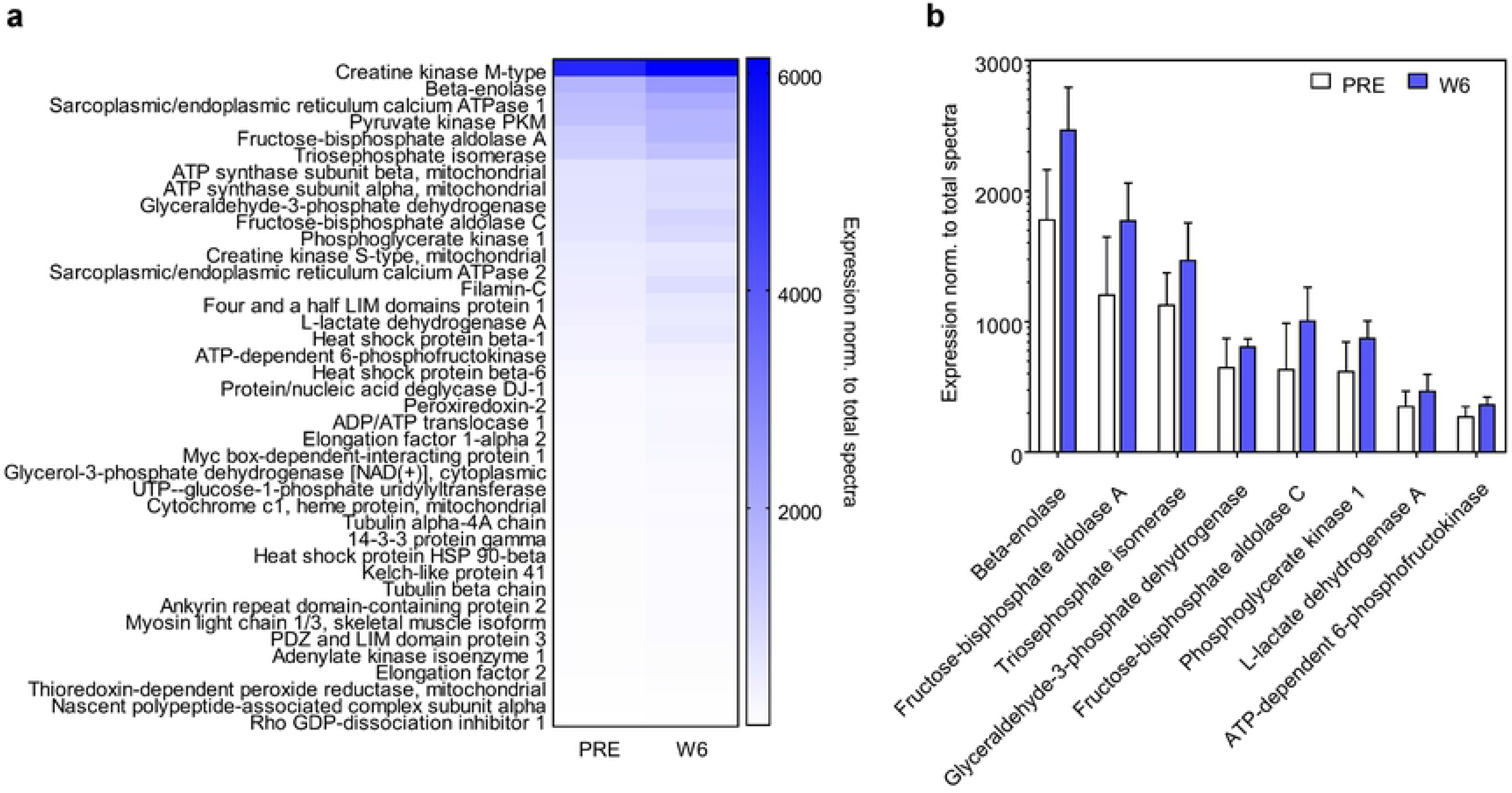
PRE and W6 sarcoplasmic fraction proteomic analysis. Legend: Proteomics analysis was performed by LC-MS/MS on n=12 participants at PRE and W6. Out of the 157 sarcoplasmic proteins detected using proteomics analysis, 40 proteins were significantly upregulated from PRE to W6 (p<0.05; panel a). KEGG pathway analysis indicated that only one pathway termed glycolysis/gluconeogenesis (8 up-regulated enzymes) was significantly up-regulated from PRE to W6 (panel b). Data in panel b are presented as mean ± standard deviation values.

### W7 VL biopsy variables and serum CK activity levels in a subset of fCSA responders

Figure 5 demonstrates data from the 7 fCSA responders that refrained from training 8 days following the last training bout. The repeated measures ANOVA p-value for mean fCSA was 0.152 (Figure 5a). Given that the n-size was limited for these data we performed a forced post hoc and confirmed that fCSA increased from PRE to W6 in these 7 fCSA responders (p=0.006). However, W7 fCSA was not significantly different from PRE (p=0.326) or W6 (p=0.515). Sarcoplasmic protein content significantly increased from PRE to W6 (p=0.003), and remained elevated at PRE versus W7 (p=0.011) (Figure 5b). Myosin and actin concentrations were significantly lower at W6 and W7 versus PRE (p<0.05 for both targets), and trended downward from W6 to W7 (p<0.10 for both targets) (Figure 5c and d). CS activity decreased from PRE to W6 and W7 (p<0.05 at both levels of time) (Figure 5e). Interestingly, while muscle fluid content did not increase from PRE to W6, this metric trended downward from W6 to W7 (p=0.056; Figure 5f). There were no significant changes in muscle glycogen concentrations (Figure 5g), serum CK activity (Figure 5h) or muscle 20S proteasome activity (Figure 5i).

**Figure 5.**
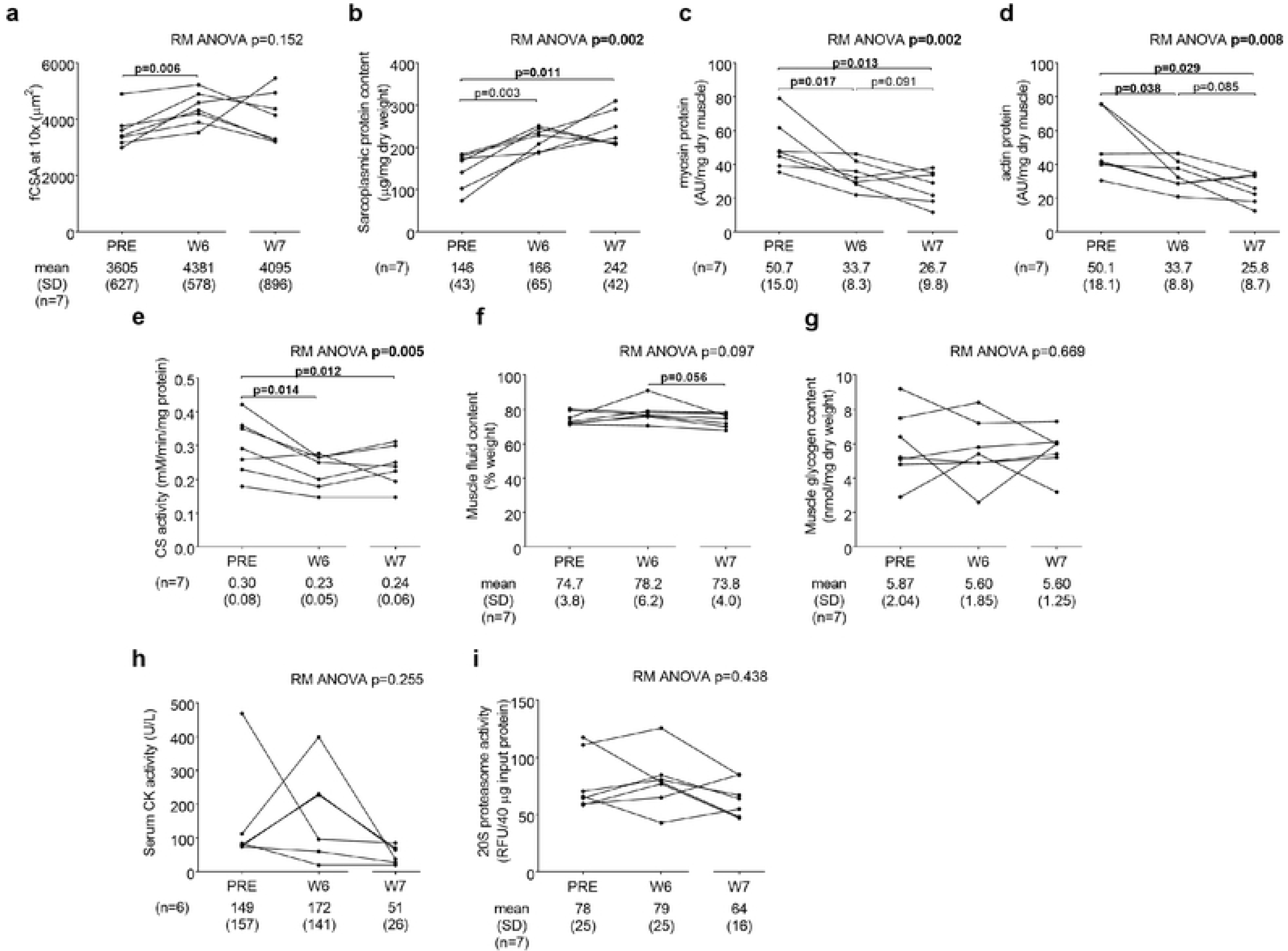
Actin and myosin decrements and concomitant increases in sarcoplasmic protein concentrations persist after one week of detraining. Legend: All line graph data show individual responses with mean ± standard deviation (SD) values below each time point. fCSA significantly increased from PRE to W6, although W7 fCSA was not different from PRE or W6 (panel a). Sarcoplasmic protein content significantly increased from PRE to W6, and remained elevated at PRE versus W7 (panel b). Actin and myosin concentrations were significantly lower at W6 and W7 versus PRE, and trended downward from W6 to W7 (panels c and d). CS activity decreased from PRE to W6 and W7 (panel e). While muscle fluid content did not increase from PRE to W6, this metric trended downward from W6 to W7 (panel f). There were no significant changes in muscle glycogen concentrations (panel g), serum CK activity (panel h) or muscle 20S proteasome activity (panel i).

## DISCUSSION

There is a widespread consensus that resistance training-induced skeletal muscle hypertrophy involves mean fCSA increases due to proportional increases in contractile proteins. In this regard, several reports have implied that the addition of sarcomeres in parallel and/or the addition of new myofibrils are largely responsible for muscle fiber hypertrophy [33-35]. This hypothesis seems logical given that elegant tracer studies have consistently demonstrated that high-volume resistance exercise increases MyoPS days following or weeks during the training stimulus. Notwithstanding, and as discussed earlier, several studies suggest skeletal muscle myofibril protein density may actually decrease following weeks to years of higher volume resistance training in humans [16-18]. Furthermore, a recent study suggests that lower specific tensions are evident in single fibers isolated from bodybuilders with significantly larger fCSAs relative to control and power athlete fibers [36]. These authors explicitly stated “*hypertrophy has a detrimental effect on specific tension*”, and noted that myofibril dilution through higher volume body building-style training may have been a driving factor for their observations.

In the current study, an intriguing observation was that actin and myosin concentrations decreased in lieu of fCSA increases over 6 weeks of high-volume resistance training in previously-trained participants. Phalloidin staining verified that actin content per fiber did not change in lieu of hypertrophy, and suggests that actin filament density decreased. Additionally, a dilution effect in mitochondrial content was also observed through observed decrements in muscle CS activity. It is possible that the extraordinarily high training volumes may have accelerated muscle proteolysis or muscle damage leading to a loss of actin, myosin and mitochondrial proteins. While proteolysis rates were not determined at each time point, muscle 20S proteasome activity and polyubiquinated protein levels (surrogates of proteolysis) were not significantly different from baseline at the time points subsequent samples were collected. However, this does not eliminate the possibility proteolytic rates were transiently increased at time points prior to or following inquiry and the underlying mechanisms of our findings warrant further research. Moreover, there was a numerical increase in sarcoplasmic protein concentrations from PRE to W6 that trended towards significance (p=0.065; data not shown due to RM ANOVA p>0.100). Collectively, these data suggest that: a) sarcoplasmic expansion (i.e., sarcoplasmic hypertrophy) was the primary mode through which muscle hypertrophy occurred from PRE to W6, and b) this effect seemed to persist up to 8 days following the last training bout in the subset of participants that were analyzed at the W7 time point.

Our interpretation of data presented in Figures 1 and 2 was that sarcoplasmic expansion occurred from PRE to W6. Thus, we elected to perform a proteomic analytical approach of the sarcoplasmic protein fractions at these time points. Bioinformatics suggested that there was a significant up-regulation of numerous enzymes involved in glycolysis from PRE to W6. Aside from this pathway, it is also notable that several enzymes involved in ATP generation were also upregulated including two creatine kinase isoforms (cytosolic and mitochondrial), two ATP synthase subunits (alpha and beta), ADP/ATP translocase, and adenylate kinase. In an attempt to logically harmonize our findings with existing knowledge pertaining to muscle physiology, our interpretations of these observations are as follows: a) high-volume resistance training may promote sarcoplasmic expansion to spatially prime cells for the expansion of the myofibril pool, b) an increase in the expression of enzymes involved in ATP re-synthesis and metabolism is sensible considering that myofibrillar protein expansion requires large amounts of ATP due to the energetic costs of synthesizing and assimilating myofibrillar proteins, and c) an upregulation in sarcoplasmic proteins related to ATP generation and excitation-contraction coupling may also occur in lieu of sarcoplasmic expansion due to the metabolic demands of high-volume training as well as the energetic demands of myofibrillar protein synthesis and assimilation.

Considering these findings, we posit that a reappraisal of dogma regarding high-volume resistance training-induced hypertrophy is warranted. In this regard, some have argued that the utilization of high-volume training to volitional fatigue promotes skeletal muscle hypertrophy to a similar degree when compared to lower volume/higher load training [37]. This argument is supported by numerous studies which have determined that both training modalities similarly increase surrogates of skeletal muscle hypertrophy [1, 38-40]. However, these conclusions have primarily been drawn through the utilization of ultrasound to assess muscle thickness or fCSA assessments using histological staining techniques, and none of these studies have examined if the stoichiometric relationship between myofibrillar and sarcoplasmic protein concentrations are preserved relative to pre-training levels. Stated differently, it is unclear in these studies the specific mode through which hypertrophy occurred (e.g., sarcoplasmic, myofibrillar, or connective tissue) and what the functional outcomes of these findings may be. It is also notable that higher load training consistently increases strength compared to lower load training [1, 38-40]. While speculative, it is possible that higher load training (e.g., 3-5 RM lifting) proportionally increases myofibrillar protein levels and fCSA, and this mode of ultrastructural hypertrophy promotes optimal strength gains. Conversely, higher volume/lower load training (e.g., 8-12+ RM lifting) may increase fCSA predominantly through sarcoplasmic expansion and metabolic conditioning, thus leading to sub-optimal strength gains. However, it stands to reason that a combination or specific sequence of the two training emphases (e.g., light-load, high-volume vs. heavy-load, low-volume) may produce superior strength and hypertrophy gains compared to the chronic completion of only one or the other (e.g., training periodization). Indeed, as posited by Stone et al. [41] decades ago, it is possible, if not likely, that higher-volume/lower-load training resulting in sarcoplasmic expansion and metabolic conditioning preceding lower-volume/higher-load training can produce more favorable strength outcomes. While these hypotheses are provocative, they are indeed speculative given that the current study lacks a high-load training group and research assessing cellular adaptations comparing different training styles is lacking. Therefore, studies examining the cellular adaptations to high-load/low-volume versus low-load/high-volume resistance training are warranted.

The implemented training volume was extraordinarily high and the W6 biopsy was collected 24 hours following the last training bout. Therefore, it is plausible that training-induced muscle damage and edema may have influenced many of the dependent variables reported herein. However, much of our data discounts edema being a confounding variable which lead to a decrease in actin and myosin concentrations. First, tissue fluid levels remained unaltered from PRE to W6, and had edema been a confounder then it is plausible that fluid levels would have increased. Additionally, multiple studies discussed above have reported that higher volume resistance training promotes sarcoplasmic expansion and, as noted earlier, our findings are in agreement with these reports. High-volume training-induced sarcoplasmic hypertrophy is also supported by a previous study [5] where the authors examined how 90 days of higher volume resistance training (i.e., up to 12 RM lifts) followed by 90 days of detraining affected mean fCSA. Following 90 days of training, fCSA increased 16% from pre-training values. Three days of detraining (which the authors termed ‘D3’) numerically increased fCSA 3% more from the 90-day point (i.e., the last day of training) and 19% from pre-training. However, fCSA decreased back to pre-training values after only 7 additional days of detraining following D3. In relation to the current study, the W7 biopsy data obtained herein provides very compelling evidence supporting the role of sarcoplasmic hypertrophy in facilitating the observed fCSA increases given that myosin and actin concentrations continued to decrease from W6 to W7 with a concomitant increase in sarcoplasmic protein.

### Experimental limitations and future directions

As with all studies that interrogate human skeletal muscle specimens, an experimental limitation to the current study is the extrapolation of findings from relatively small biopsy samples. Another limitation to the current study is that only college-aged males were examined. Thus, our findings may not represent the physiology that occurs in females or older populations. Due to financial constraints and the nature of the research question, the low fCSA responders were not extensively analyzed relative to the higher fCSA responders. However, in light of the −768±754 μm^2^ fCSA decrease that occurred in these 16 participants, it is notable that: a) similar to high fCSA responders, sarcoplasmic protein content significantly increased from PRE to W3 (p=0.022) and trended upward from PRE to W6 (p=0.069) [mean±SD values (μg/mg dry weight): PRE=165±75, W3=203±63, W6=214±48], b) similar to high fCSA responders, myosin protein content decreased from PRE to W3 (p=0.040) and PRE to W6 (p=0.023) [mean±SD values (AU/mg dry weight): PRE=66.6±48.4, W3=46.7±27.3, W6=28.5±16.3], and c) actin protein content decreased from PRE to W3 (p=0.034) and PRE to W6 (p=0.020) [mean±SD values (AU/mg dry weight): PRE=62.1±42.5, W3=42.7±23.1, W6=26.8±13.6]. Thus, it seems likely that a disproportionate increase in sarcoplasmic volume relative to contractile protein content also occurred in low fCSA responders, but these alterations were not sufficient enough to increase fCSA compared to baseline levels, and may be considered to indicate preparatory remodeling rather than hypertrophy. Additionally, we did not perform W3, W6, or W7 strength testing given that the primary intent of the training paradigm was to elicit skeletal muscle hypertrophy and not strength. However, in retrospect, it would have been interesting to observe if strength changes plateaued or even decreased at these time points relative to PRE, and if these changes were associated with decrements in actin and myosin protein concentrations. Finally, it remains to be determined what other myofibrillar proteins (e.g., nebulin, titin) may be expressed to differing degrees in response to various training interventions beyond myosin and actin, which are considered to occupy the majority of the myofibril, and what specific training stimuli instigate the expression of specific myofibrillar proteins.

## CONCLUSIONS

These data challenge current dogma suggesting fCSA increases during high-volume resistance training are primarily driven through increases contractile protein content. Instead, we interpret these data to suggest sarcoplasmic hypertrophy is largely responsible for short-term fCSA increases, and this effect can persist up to 8 days following the cessation of training. Our proteomic data suggest that sarcoplasmic proteins upregulated during higher volume training are involved with glycolysis as well as other processes that generate ATP. Future research is needed to determine the significance of sarcoplasmic hypertrophy during high-volume resistance training, or whether different training paradigms affect this process.

## ACKNOWLEDGEMENTS

The authors have declared that no competing interests exist. Funding for subject compensation and certain assays was provided through a contract awarded to KCY and MDR by Impedimed, Inc. Funding for proteomic analysis was provided through a Grant in Aid to MDR and VI from Florida A&M University. Additional funding was provided as a gift in kind to CTH and MDR through Renaissance Periodization by Dr. Mike Israetel and Nick Shaw. The authors would like to thank the subjects for taking the time and effort in completing this grueling training and biopsy protocol.

## SUPPORTINING INFORMATION

The datasets used and/or analyzed during the current study are available from the corresponding author upon reasonable request.

